# A pre-trained large generative model for translating single-cell transcriptome to proteome

**DOI:** 10.1101/2023.07.04.547619

**Authors:** Linjing Liu, Wei Li, Ka-Chun Wong, Fan Yang, Jianhua Yao

## Abstract

Proteins are crucial for life, and measuring their abundance at the single-cell level can facilitate a high-resolution understanding of biological mechanisms in cellular processes and disease progression. However, current single-cell proteomic technologies face challenges such as limited coverage, throughput, and sensitivity, as well as batch effects, high costs, and stringent experimental operations. Drawing inspiration from the translation procedure of both natural language processing (NLP) and the genetic central dogma, we propose a pre-trained, large generative model named scTranslator (single-cell translator). scTranslator is align-free and capable of generating multi-omics data by inferring the missing single-cell proteome based on the transcriptome. Systematic benchmarking confirms the accuracy, stability, and flexibility of scTranslator across various quantification techniques, cell types, and conditions. Furthermore, scTranslator has demonstrated its superiority in assisting various downstream analyses and applications, including gene/protein interaction inference, gene pseudo-knockout, cell clustering, batch correction, and cell origin recognition on pan-cancer data.

## 1 Introduction

Since the advent of single-cell transcriptome sequencing (scRNA-seq) technology [1], research on biological processes has entered the era of single-cell analysis, mainly based on transcriptomics [2]. Although scRNA-seq methods have provided information about the RNA landscape in many biological systems and demonstrated high clinical relevance, their readouts as proxies for protein levels are limited [2–4]. As the main drivers of cellular processes [5], protein level is essential for capturing the molecular mechanisms in cell differentiation and fate determination, cellular signaling pathways, and disease progression. However, the technical limitations in single-cell proteomics hinder the widespread generation and analysis of the corresponding data. Therefore, inferring protein abundance through computational methods at the single-cell level is a valuable and complementary approach.

At present, single-cell proteomics techniques are based on next-generation sequencing (NGS) or mass spectrometry (MS). The NGS-based techniques rely on well-validated affinity reagents and has limitations in the number of target proteins [6], typically ranging from a dozen to several hundred in actual experiments. This is insufficient given that a single cell can express over 10,000 different proteins at varying levels [7]. Besides, current available data generated by NGS are constrained to surface proteins. Although MS-based techniques provide an opportunity to analyze thousands of proteins and post-translational modifications, this technique is prone to both experimental and computational artifacts, which may ultimately compromise the quality of data [8]. Moreover, single-cell proteomic technologies are confronted with shared obstacles that include batch effects, high cost, and stringent experimental operations [6, 8].

In this regard, the development of deep learning models to predict proteome abundance could provide a potential solution. Notably, cTP-net [9] and sciPENN [10] are pioneering works in this field. However, there are still some limitations: (a) model scalability, where the two methods above are unable to utilize genes or predict proteins that have not appeared in the training set, limiting the potential of the model in unknown areas or even new datasets; (b) model flexibility, where both cTP-net and sciPENN require feature alignment before applied on new datasets; (c) prediction only about cell surface protein abundance, which is due to the limitations of single-cell sequencing technologies and the lack of pre-training on datasets involving intracellular proteins; and (d) limited downstream application, both methods only focused on the common tasks for single cells such as annotation and clustering, lacking guidance for clinical applications.

According to the central dogma of molecular biology, RNA sequences are translated into amino acid sequences that form proteins. The translation process from RNA to protein shares remarkable similarities with the task of machine translation in NLP, which involves the use of algorithms to automatically translate one language into another. The machine translation has undergone a transformative change with the advent of pre-trained language models. Inspired by the translation process in NLP and the central dogma, we propose scTranslator as the first pre-trained, context-aware, and align-free large generative model for generating multi-omics data by translating single-cell transcriptome to proteome. In this work, we collected an extensive corpus of RNA and protein from public bulk and single-cell datasets, encompassing 31 different cancer types, 18,227 patient-level samples, 239,797 cell-level samples, and a total of 76 datasets. Through the general knowledge acquired from substantial corpus of RNA and protein pairs during pre-training stage, scTranslator has the potential to generate corresponding proteome data for transcriptome, encompassing not only surface proteins but also intracellular proteins. By innovatively introducing the re-indexed Gene Positional Encoding (GPE) module into Transformer [11], scTranslator can infer any protein with arbitrary length determined by the user’s query, as the GPE module has comprehensive coverage of all gene IDs and reserves another 10,000 positions for new findings. Moreover, the GPE module links gene ID of RNA or protein with its corresponding expression level, enabling scTranslator to be an align-free model. Systematic benchmarks and case studies demonstrate that scTranslator is accurate, flexible, and scalable for generating paired protein abundance with solid interpretability mechanism demonstration.

As a pre-trained large generative model, scTranslator exhibits exceptional potential across a wide range of downstream applications. First, we employed the attention matrix of scTranslator to investigate the regulatory or interaction relationships of gene-gene, gene-protein, and protein-protein. Second, we leveraged the flexibility of scTranslator to explore the effects of specific gene on over 10,000 proteins by pseudo-knockout. Third, the proteome data generated by scTranslator achieved superior performance on cell clustering and batch correction. Finally, we demonstrated that the multi-omics data generated by scTranslator, which includes both surface and intracellular proteins, can aid in the identification of cells from tumor tissues in a pan-cancer dataset. Based on these findings, we anticipate that scTranslator could improve understanding of regulatory and interaction relationships at a high-resolution level and facilitate the research of single-cell multi-omic analysis.

## 2 Overview of scTranslator and its downstream applications

Transformer-based [11] models are powerful architectures in both natural language processing and computer vision [12–16]. The prevailing approach is to pre-train on large text corpus or image datasets, and then fine-tune on downstream task-specific data sets [15, 17–19]. As pre-training has been shown to be effective for improving the performance of many tasks [20–23], we develop scTranslator following a similar paradigm with the dominant approach. An overview of scTranslator and its application on pan-cancer data is presented in Fig. 1.

**Fig. 1.**
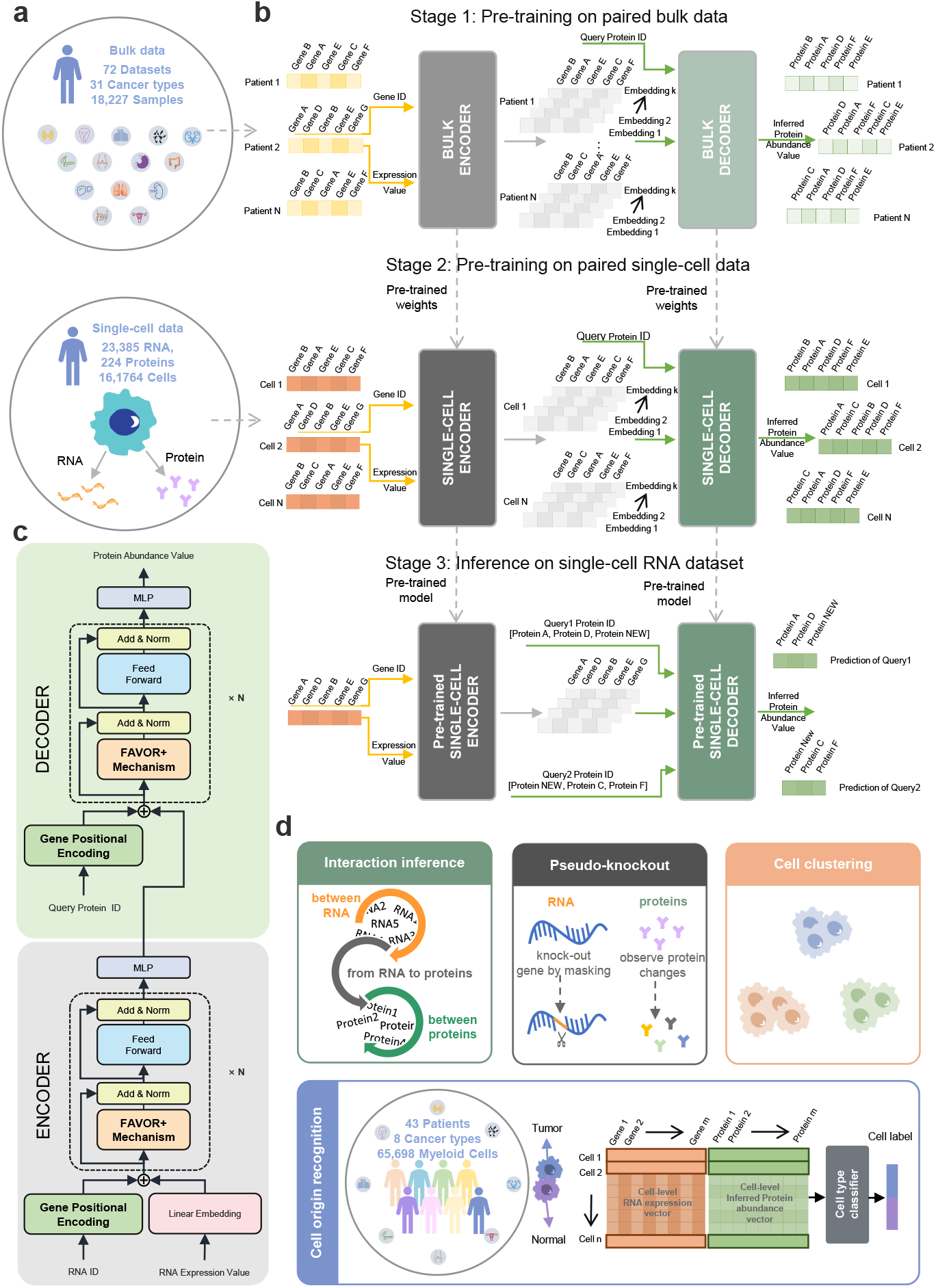
Overview of scTranslator and its downstream application. **a,** A summary of bulk and single-cell data for the pre-training stage. **b,** The three stages of training and utilizing scTranslator. Stage 1: Pre-training on paired bulk dataset; Stage 2: Pre-training on paired single-cell dataset, Stage 3: Inferring on single-cell RNA dataset with or without fine-tuning. We adopted MSE loss to train the model. **c,** The encoder-decoder architecture of scTranslator with tailored design for transcriptome and proteome data. **d,** The downstream applications of scTranslator. The top panel shows applications on interaction inference, pseudo-knockout, and cell clustering. The bottom panel is a summary of single-cell pan-cancer data illustrates the use of generated multi-omics data for cell origin recognition.

Datasets used for pre-training are depicted in Fig. 1a. We first collected all available paired RNA and protein data from the Cancer Genome Atlas, Memorial Sloan Kettering Cancer Center, the Broad Institute and Novartis, and Clinical Proteomic Tumor Analysis Consortium, named paired bulk data. The bulk data consists of 72 datasets, covering 31 cancer types and involving 18,227 patient-level samples. Given the maturity of bulk sequencing for joint profiling of transcriptome and proteome, the measured proteins comprise not only intracellular proteins but also surface proteins. The second type of pre-training data is paired single-cell data collected from reference ([24]), which is the largest available dataset in terms of protein and cell amount.

There are three stages in our framework as illustrated in Fig. 1b. In stage 1, scTranslator is trained on paired bulk data and each sample comes from a patient. In stage 2, the pre-trained scTranslator continues training on paired single-cell data and each sample comes from a cell. In stage 3, we can use the pre-trained scTranslator from the previous two stages to infer protein abundance on single cell RNA datasets with or without fine-tuning. Throughout all training stages of scTranslator, we utilized the mean square error (MSE) between the actual and predicted proteins as the loss function. Details of scTranslator and pre-training strategy are in the Methods section.

As presented in Fig. 1c, scTranslator utilizes the encoder-decoder architecture, similar to Transformer, and integrates customized design for transcriptome and proteome data. Notable modifications include the re-indexed GPE, Fast Attention Via positive Orthogonal Random features approach (FAVOR+) [25], and one-forward generation of decoder. scTranslator accepts RNA expression value and corresponding gene ID as input. The expression value is projected to a pre-set dimensions vector by a learnable embedding layer. Position encodings are added to the expression embedding to distinguish information from different genes. We developed re-indexed GPE as a learnable position encoding that converts NCBI Gene ID and new findings into position vectors with the same dimension as expression embeddings. For scTranslator decoder, the position encodings are embeddings of protein ID queried by any user with arbitrary length. Since our sequence length far exceeds the sentence length in NLP, to reduce the time and space complexity, scTranslator adopts FAVOR+ as the multi-head attention mechanism in both the encoder and decoder. FAVOR+ allows scTranslator to capture long-range dependencies and context information more effectively. In addition, unlike the traditional Transformer, scTranslator does not employ an auto-regressive decoder, which predicts the requested protein abundance in a step-by-step manner. The generative style decoder of scTranslator predicts with one forward operation, thereby improving the inference efficiency of long-sequence predictions [26].

In this study, we performed a series of downstream applications (Fig. 1d) including interaction inference, gene pseudo-knockout, cell clustering. Specifically, we applied scTranslator to a pan-cancer dataset, where only scRNA-seq data is accessible, to infer the absent protein abundance. The pan-cancer dataset contains myeloid cells from 43 patients with eight types of cancer. We utilized both scRNA-seq data and predicted proteomic data to distinguish cells derived from tumor versus normal tissue, and found the predicted proteins could assist in identifying cells from tumor tissue.

## 3 Results

### 3.1 scTranslator enables precise prediction on both bulk and single-cell data

As scTranslator has undergone two stages of pre-training, it is imperative to assess the performance of this extensive and intricate pre-training task to establish the dependability of the pre-trained model before advancing to downstream tasks. We pre-trained the model on 90% of the data and evaluated the effectiveness of the pre-training using the remaining 10% of the data as the test set. At the end of each pre-training stage, we conducted sample-wise evaluations by comparing predicted values with ground truth for each sample (patient or cell). From Fig. 2a, scTranslator has extremely close performance on training data and test data, indicating that it generalizes well on unseen data. Furthermore, it achieves superior performance in terms of both cosine similarity and MSE, regardless of the type of sequencing technique. In particular, scTranslator attains a cosine similarity of over 0.98 and an MSE of less than 6 × 10*^−^*^3^ on both the training and test sets obtained from bulk sequencing. For single-cell sequencing data, scTranslator yields a cosine similarity of over 0.93 and an MSE of less than 2 × 10*^−^*^3^ on both datasets. The results suggest that scTranslator exhibits good generalization on previously unseen data, attesting to the model’s dependability and capacity to grasp the underlying RNA-protein relationship. Overall, these outcomes demonstrate the success of scTranslator’s pre-training stage and underscore its potential for downstream applications.

**Fig. 2.**
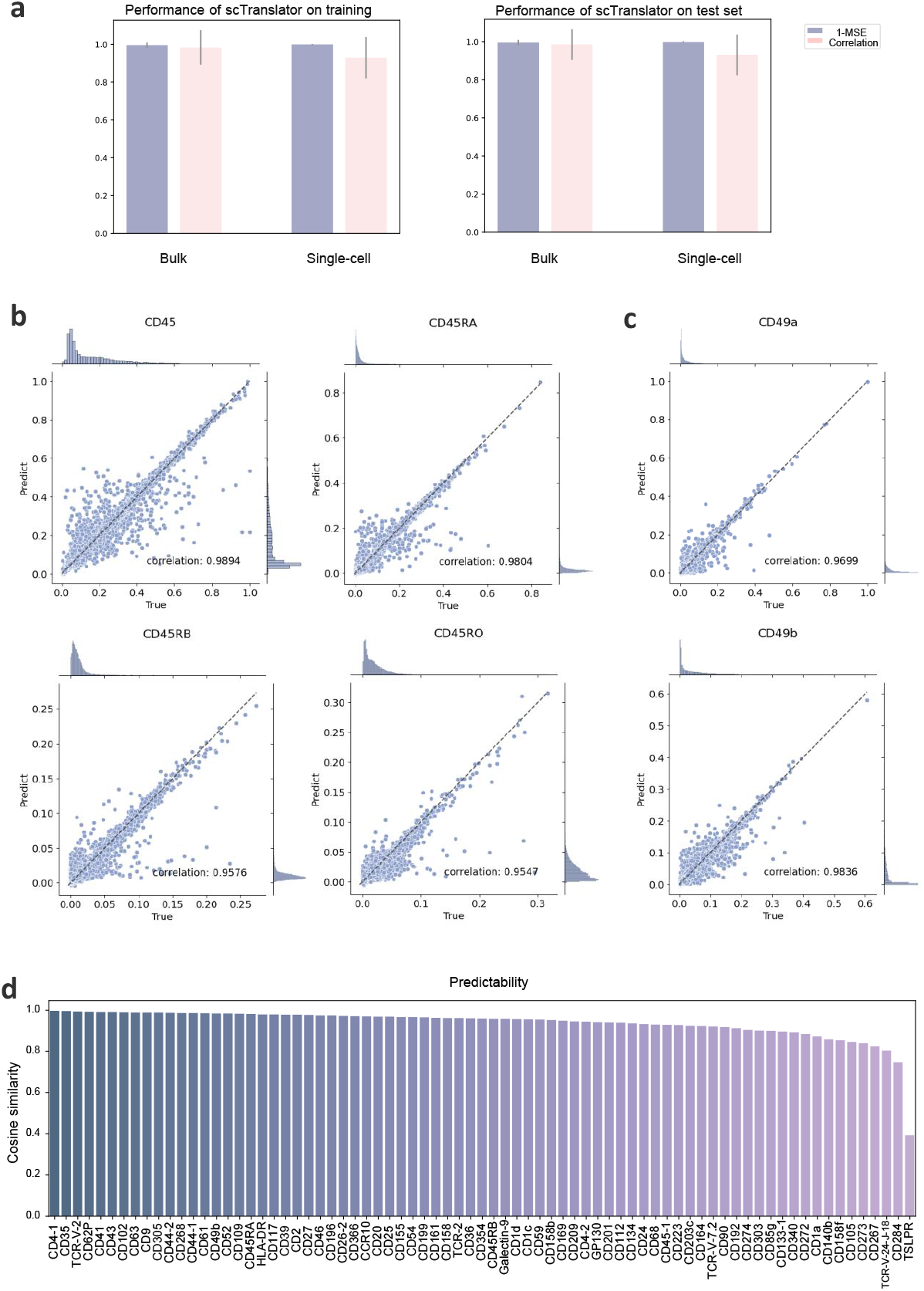
Overall performance of scTranslator. **a,** Performance of scTranslator evaluated at sample-wise level by computing MSE and cosine similarity between the true and predicted values for each sample (patient or cell). The left panel and right panel are reported on the training set and test set, respectively. The error bars indicate mean *±* s.d across all samples. **b,** Joint plots of ground truth and prediction, for CD45 and its isoforms. For each subplot, the distribution histograms of the predicted and true value are displayed on the border, and the middle part is their scatter plot. **c,** Joint plots of ground truth and prediction for protein CD49a and CD49b. More joint plots of individual proteins and their family members are shown in Extended Data Fig. A1. **d,** The predictability of each protein evaluated based on cosine similarity. Higher cosine similarity indicates better predictability.

Notably, single-cell datasets contain both proteins and their isoforms, which are produced by alternative splicing and exhibit similar function. This coexistence poses a challenge to the model’s capacity to accurately predict the abundance of diverse isoforms. However, as depicted in Fig. 2b, the joint plots of actual and predicted values for CD45, CD45RA, CD45RB, and CD45RO demonstrate the ability of scTranslator to precisely predict the abundance of both the parent protein and its isoforms. Fig. 2c are joint plots from CD49 protein family member and more joint plots of individual proteins are provided in Extended Data Fig. A1. The comprehensive performance of scTranslator on all proteins (Fig. 2a), coupled with the meticulous evaluation of concordance between predicted and observed values on a single protein (Fig. 2b,c and Extended Data Fig. A1), indicates that our model not only exhibits accuracy in protein prediction at a global level but also manifests the commendable ability to depict distribution at an individual level.

From a biological point of view, information flows from genome to proteome, thereby realizing genotype to phenotype [27–30]. Whether RNA can be used as a substitute for protein and to what extent differences between mRNAs are reflected at the level of proteins are important in the process and have attracted lots of researches [27, 31]. Single-cell multi-omics technology allows us to simultaneously obtain the proteome and transcriptome at the single-cell level, providing a valuable opportunity to explore the above issues between RNA and proteins at the single-cell level. Here we use the proposed deep learning model to fit the relationship between mRNA and protein abundance in a non-linear and comprehensive perspective. The predictability of each protein in terms of cosine similarity is presented in Fig. 2d. Due to the canvas limitation, only one-third of the proteins are listed in the figure, while a Supplementary Table 1-1 is provided to detail protein predictability. The predictability ranges from 0.39 to 0.99 in PBMCs, most likely reflecting the differences in translation rate and protein degradation for individual proteins [27, 32]. Fig. 2d shows that a significant proportion of proteins, approximately 97%, display a correlation between prediction values and ground truth exceeding 0.8. The enrichment results (Supplementary Table 1-2) show a tendency that the proteins with higher predictability are enriched for molecular transducer activity, signaling receptor activity, and MHC protein binding. In the contrary, the proteins with lower predictability are enriched for regulation of adaptive immune response and immune effector process (Supplementary Table1-3). The above tendency must be further investigated by including more single-cell multi-omics data in more cell lines and tissues along with the popularization of single-cell multi-omics sequencing technology, so as to obtain more definite conclusions.

### 3.2 Systematic benchmarking demonstrates the remarkable performance of scTranslator

In this study, we benchmarked scTranslator against state-of-the-art (SOTA) models in protein abundance prediction task using three gold-standard paired single-cell RNA and protein datasets derived from diverse sequencing technologies or laboratory platforms. Specific details on these datasets can be found in the Method section. To ensure a grim comparison, we included the two most advanced protein abundance prediction models, cTP-net and sciPENN, for comparison. Furthermore, scTranslator is the first pre-trained model designed for single-cell protein abundance prediction. To assess the impact of the pre-training approach, we conducted a performance comparison between scTranslator with pre-training and without pre-training, which we refer to as scTranslator-scratch.

In order to provide a thorough evaluation of scTranslator’s performance, we conducted a series of experiments on these datasets. Fig. 3a illustrates three comparison experiment schemes we adopted, namely aligned, unaligned, and few-shot experiments. The aligned experiment was designed for situations where the downstream application has matched transcriptome and proteome data between the training and test sets. To demonstrate scTranslator’s exceptional capabilities as an align-free model, we employed the unaligned experiment by shuffling and randomly selecting certain genes. This results in unmatched RNA or proteins between the training and test sets, allowing us to showcase the model’s proficiency in handling un-aligned data. This aspect is crucial to its utility in downstream applications since in real scenario, as most clinical cohorts are typically sequenced using a variety of technologies, which may detect different sets of genes. In both aligned and unaligned experiments, for each dataset we utilized 90% of the data for fine-tuning and 10% for testing. To evaluate the efficacy of pre-training, a few-shot experiment was conducted, where only 20 cells were used for training and the remaining cells were used for testing. The few-shot experiment was employed to gauge the pre-trained model’s capacity to acquire knowledge from a restricted amount of data and exhibit proficiency in generalizing to novel cellular contexts. To evaluate the variability of our results and determine whether the observed effects are consistent or due to chance, we present the findings from a total of 10 experimental runs, achieved by varying the seeds to generate different splits of the dataset.

**Fig. 3.**
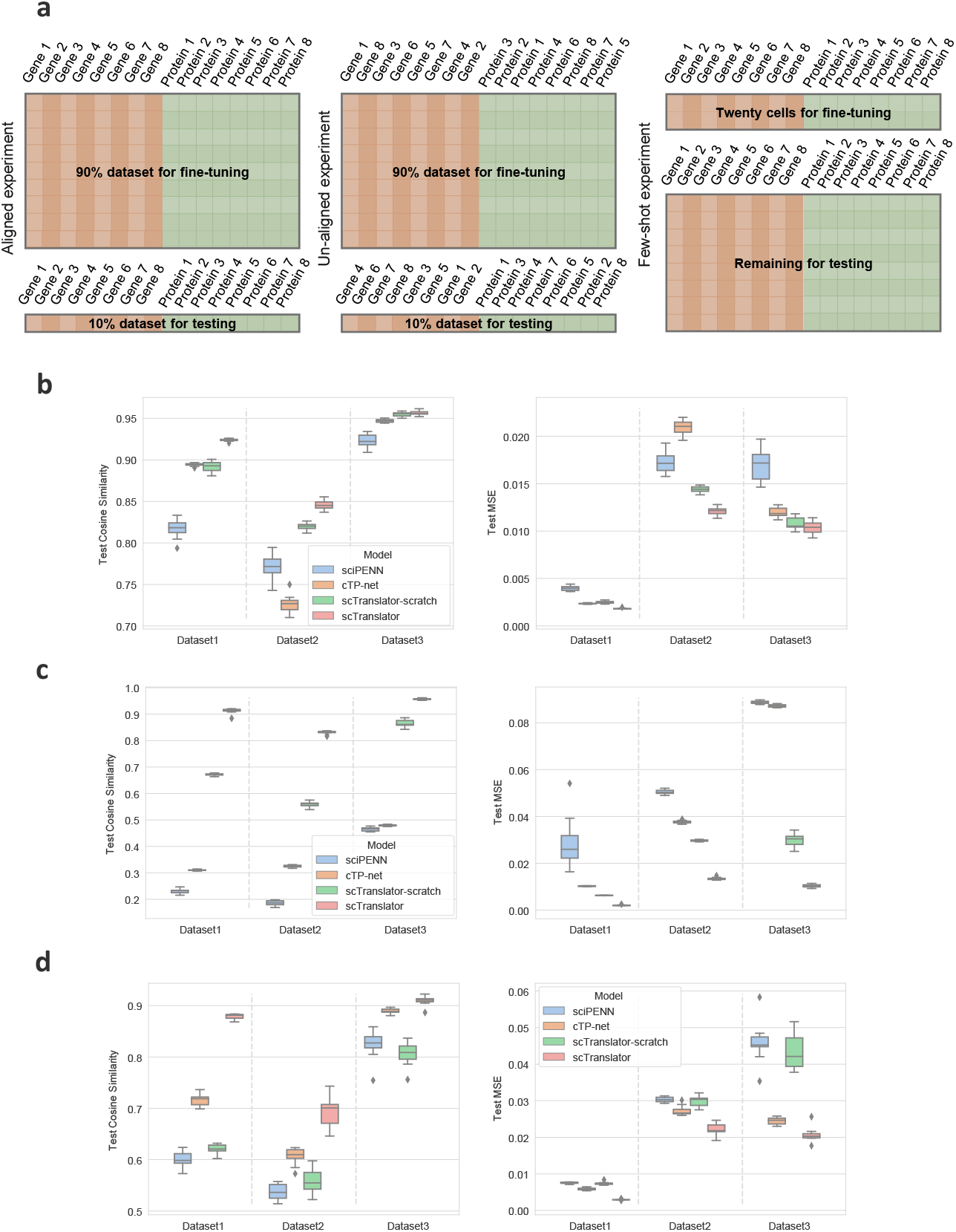
Systematic benchmarks on protein inference. **a,** The illustration of aligned, unaligned, and few-shot comparison schemes, including the dataset splitting and position align/un-align approach. **b-d,** Comparison experiment results of aligned (**b**), unaligned (**c**), and few-shot (**d**) mode. The left panel displays the cosine similarity performance of various methods on the test sets, while the right panel showcases the MSE performance. All plots are reported based on 10 runs. In each repeat, we change the random seed for different data splitting.

Fig. 3b presents comparison results in the aligned experiment, showcasing testing performance on three datasets. Further details regarding the comparison results can be found in the Supplementary Table 2. Compared to the latest SOTA sciPENN model, scTranslator surpasses it both in cosine similarity and MSE. Specifically, we observe a 10.7% increase in cosine similarity from 0.817 to 0.924 on Dataset 1, a 7.5% increase from 0.771 to 0.846 on Dataset 2, and a 3.3% increase from 0.923 to 0.956 on Dataset 3. Additionally, compared with the SOTA cTP-net model, scTranslator achieves a significant improvement in test cosine similarity on Dataset 1 and Dataset 2, with 3.0% and 11.9%, respectively. However, the performance enhancement on Dataset 3 is only about 1%. This can be attributed to the fact that Dataset 3 contains only 13 proteins, making it an easy task for all deep neural networks to learn the distribution of such a small number of proteins.

Fig. 3c depicts the outcomes of the un-aligned experiment. As sciPENN and cTP-net lack the capability to encode gene positions, the performance of both models is limited, evidenced by the cosine similarity values not surpassing 0.5 across all datasets in Fig. 3c. In contrast, scTranslator, benefiting from its GPE module, achieves significantly high cosine similarity scores of 0.911, 0.830, and 0.956 across the three test sets. These findings serve as further evidence of the effectiveness of scTranslator as an align-free model and its potential for improving protein expression inference in diverse datasets.

Fig. 3d displays the comparison outcomes of the few-shot experiment under the aligned mode, where the only difference from the aligned experiment is the extremely limited number of training samples for fine-tuning. The experimental findings indicate that scTranslator outperforms the SOTA, with a greater improvement in the few-shot experiment than in the aligned experiment. For instance, when considering cosine similarity, scTranslator exhibits a minimum increase of 2.0% over cTP-net on Dataset 3 and a maximum increase of 27.9% over sciPENN on Dataset 1. These few-shot experiment results further demonstrate the high efficiency of scTranslator in predicting protein abundance levels when transferred to new datasets, even with limited training data.

In Extended Data Fig. A2, we explored the influence of sample amount on scTranslator performance in the few-shot experiment under both aligned mode and unaligned mode. We found that the model performance benefits with the increasing of training data, however, 20 cells enable the model to achieve 80%-95% performance, compared with using the 90% dataset.

It is noteworthy to observe that, while scTranslator-scratch does not surpass scTranslator, it demonstrates competitive or superior performance compared to cTP-net and sciPENN in both aligned and unaligned experiments. This observation yields two important implications. Firstly, the superiority of scTranslator over scTranslator-scratch in all experiments suggests the effectiveness of our two-stage pre-training strategy. Secondly, the superior performance of scTranslator-scratch over the other models highlights the excellence of our model architecture in capturing the relationship between transcriptome and proteome for predicting protein abundance. However, in few-shot experiments, cTP-net beats scTranslator-scratch. This illustrates that scTranslator, as a large generative model, requires more extensive training data than cTP-net, necessitating the implementation of pre-training when constrained by the size of the downstream dataset.

From a general view, all box plots in Fig. 3 reveal that our scTranslator has the highest accuracy and the relatively narrowest distribution on most of datasets, indicating consistent and outstanding performance across different experiment runs. The rigorous series of comparison experiments conducted in this study provide compelling evidence that scTranslator is a powerful tool for protein abundance inference.

### 3.3 Integrative regulatory inference with scTranslator

In addition to providing excellent predictive capabilities for the entire proteome, scTranslator can also explore data-driven regulatory mechanisms through the inherent interpretability of the model. Based on the self-attention mechanism in scTranslator, we can observe the attention matrix from gene to protein (Fig. 4a), from gene to gene (Extended Data Fig. A3a left panel), and from protein to protein (Extended Data Fig. A3a right panel). Fig. 4a shows that some genes have larger attention scores for most of the proteins than other genes (top panel), along with differences in attention scores on various proteins (bottom two panels), and these genes may be potential dominant regulatory genes. Functional enrichment analysis showed that the genes with the top 5% sum of attention score were mainly involved in the regulation of cellular processes, cellular response to stimulus, and other mechanisms (Fig. 4b). To a certain degree, these findings are supported by the fact that all proteins measured by CITE-seq are located on the surface of the cell membrane. In turn, the regulatory genes of the proteins of interest could also be observed from the attention score. For example, Extended Data Fig. A3b lists the regulatory genes associated with TCR *γ/δ*, TCR *α/β*, CD45, and CELC2 proteins, along with their corresponding attention scores. The results (Extended Data Fig. A3c) showed that the regulatory gene list corresponding to the TCR family was relatively similar, while the gene regulatory list corresponding to the CELC2 protein was different from the TCR family (Extended Data Fig. A3c). In light of the fact that CD45 participates in TCR signaling and regulation [33, 34], the regulatory genes of CD45 show a significant overlap with TCR *γ/δ* and TCR *α/β* (Extended Data Fig. A3d).

**Fig. 4.**
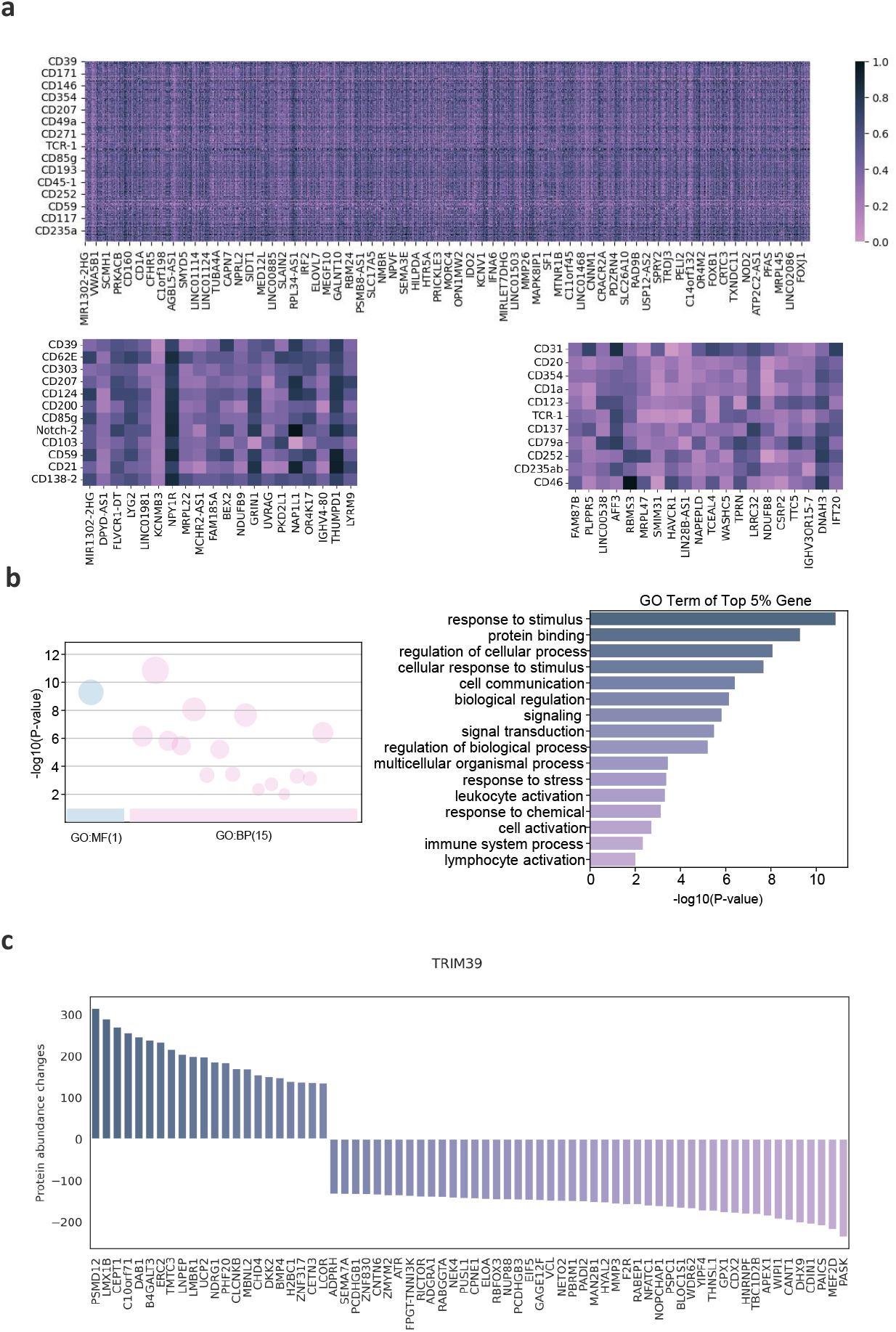
Integrative regulatory inference by model interpretability and pseudo-knockout. **a,** Heatmap of attention score matrix between RNA and protein. The top diagram provides an overview of the attention matrix, with RNA on the horizontal axis and proteins on the vertical axis. The bottom subdiagrams zoom in to show differences in attention scores between different proteins for a specific RNA of interest. **b,** Functional enrichment results based on attention scores. We computed the sum of attention scores across all proteins for each gene, and the left panel provides a summary of the number and sources of enriched GO terms for corresponding ranked genes. The right panel shows detailed Gene Ontology (GO) enrichment terms for the top 5% genes ranked by this sum. **c,** Results of TRIM39 pseudo-knockout. The vertical axis shows the sum of changes in protein abundance across all cells, and the horizontal axis shows the affected proteins at the top of the order.

Besides the attention mechanism, the model can also perform pseudo-knockout experiments for functional gene discovery in a computational way. Specifically, we can mask a specific gene from RNA expression matrix, and then observe the changes in the model’s predicted values for all proteins including surface and intracellular proteins. This process is similar to the biological CRISPR/Cas9 knockout experiment, but greatly shortens the time and experimental resources required for equivalent biological experiments. From this perspective, pseudo-knockout observations are provided for research groups that have difficulties to conduct wet experiments, providing a rich resource for new hypotheses and novel discoveries. For example, pseudo-knockout of the TRIM39 gene has a broad range of effects on a series of downstream proteins, which is consistent with the biological role of this E3 ubiquitin-protein ligase (Fig. 4c) [35]. We also pseudo-knocked out some key genes that have been reported for tumor progenesis and obtained data-driven analysis results, which provided valuable resources to be mined for tumor research (Extended Data Fig. A4). Taking AKT1 as an example, its regulatory proteins include H1-0 and ANTXR1, which are consistent with the reported biological conclusions [36, 37], proving the application potential of the model analysis results in tumor research.

### 3.4 Pseudo-protein quantification based on scTranslator assists downstream application

The ability to identify cell types accurately is crucial for understanding the cellular basis of complex biological processes. In this study, we investigated whether pseudo-protein quantification utilizing scTranslator aids in the recognition of cell types. We applied the pre-trained scTranslator to directly infer protein expression in more than 1,6000 human peripheral blood mononuclear cells (PBMCs) of Dataset 1. The uniform manifold approximation and projection (UMAP) plots, presented in Fig. 5a-d, illustrate the predicted protein abundance, raw protein data, and raw RNA data for seven cell types, namely B cells (marked by CD19), CD4 T cells (marked by CD4), CD8 T cells (marked by CD8a), other T cells (marked by CD195), dendritic cells (marked by CD1c and CD304), monocytes (marked by CD11b), and natural killer cells (marked by CD224). Other markers that have been reported are listed in the Supplementary Table 3. Our findings indicate that, in line with observations from CITE-seq studies [38], predicted protein levels, similar to the actual protein values, differ substantially from RNA expression level of the corresponding gene, exhibiting greater contrast across cell types and greater uniformity within each cell type. Moreover, we identified potential novel cell markers that have not been previously reported, as visualized in Extended Data Fig. A5 and Extended Data Fig. A6a.

**Fig. 5.**
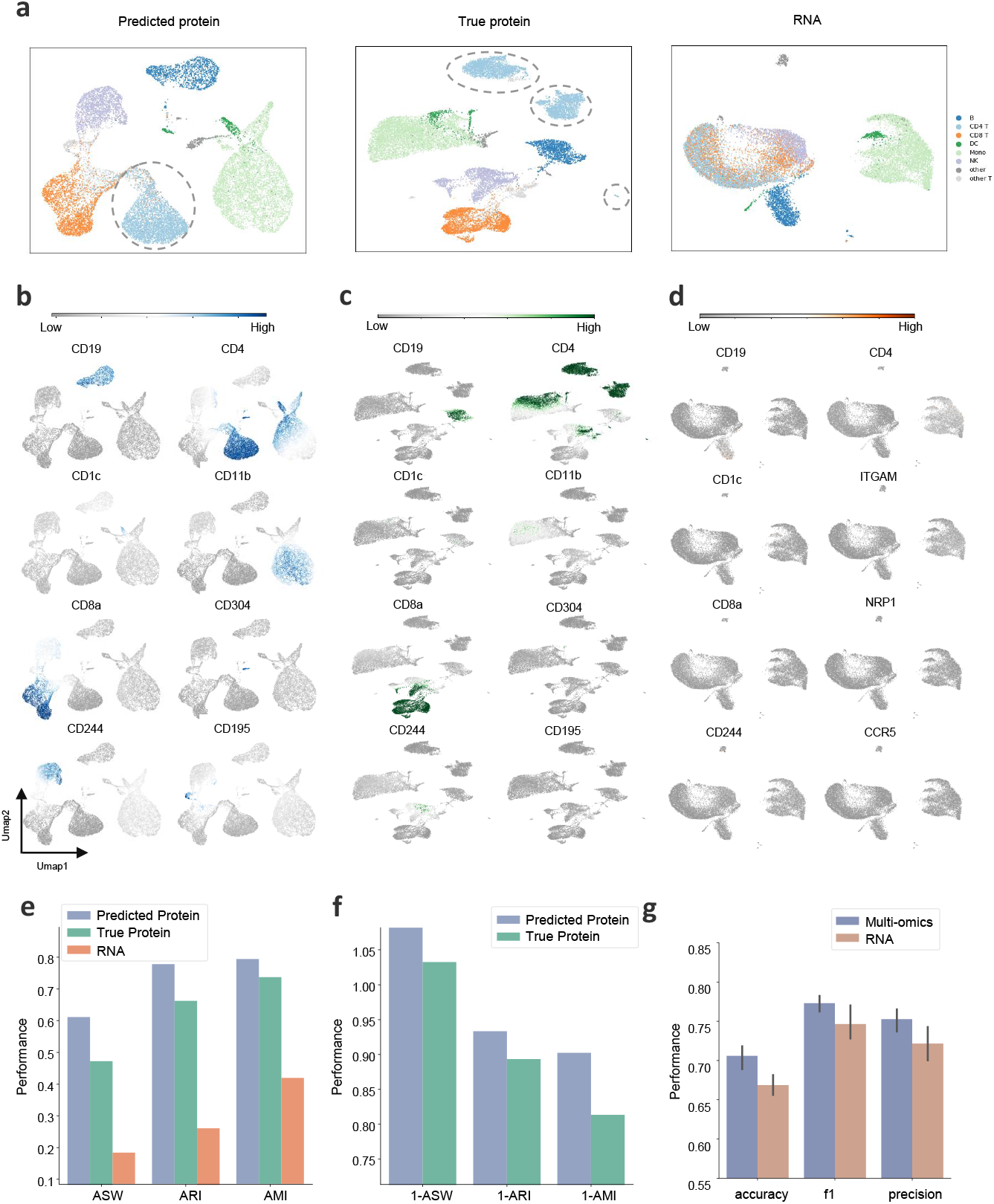
Exploration of cell heterogeneity in Seurat v4 PBMCs and cell origin in pan-cancer data. **a,** Visualization of Dataset 1 colored by cell type. The left, middle, and right are plots of predicted protein, true protein, and raw RNA, respectively. The dotted circles signify the presence of a batch effect in the raw data, which is effectively eliminated by the predicted values generated by scTranslator. Detailed batch information is shown in Extended Data Fig. A6b. **b-d** UMAP plots colored by the level of eight reported markers for different cell types. The UMAP visualized inferred protein (**b**), true protein (**c**), and raw RNA (**d**). Novel markers found in this study are visualized in Extended Data Fig. A5 and Extended Data Fig. A6a. **e,** Cell clustering performance evaluated by ARI, AMI, ASW, and HS. **f,** Batch correction performance evaluated by 1-ARI, 1-AMI, 1-ASW, and 1-HS. **g,** Cell origin recognition performance evaluated by accuracy, f1, and recall.

Remarkably, scTranslator appears to have acquired a general pattern of the interplay between RNA and protein from its pre-training stage, which endows our model to remain unaffected by batch diversity while simultaneously preserving crucial information related to cellular heterogeneity. In the middle panel of Fig. 5a, three distinct clusters of CD4 T cells are observed in the visualization of true protein. Batch effects were found to influence these clusters, as shown in Extended Data Fig. A6b, which depicts batch information from eight donors. Specifically, the first cluster was predominantly composed of cells from donors 1-4, the second cluster comprised cells from donors 5-8, while the third cluster included some CD4 T cells from donor 7. In contrast, the left panel of Fig. 5a demonstrates that the predicted proteins effectively correct batch effects, resulting in a more uniform clustering of CD4 T cells and the formation of a single cluster.

To quantitatively evaluate whether scTranslator eliminates the batch effect while preserving the heterogeneity information between cell types, we compared cell clustering performance across predicted proteins, true proteins, and RNA data, as well as the batch correction performance between predicted protein and true protein. For cell clustering, we employed K-means clustering with the given cell types as ground truth and assessed the results using adjusted Rand index (ARI), adjusted mutual information (AMI), and adjusted silhouette width (ASW). A higher value for these metrics indicates a better correspondence with the true cell type. The results in Fig. 5e demonstrate that pseudo-protein quantification based on scTranslator achieved accurate clustering with top-ranking performance on all metrics, outperforming true protein and RNA data. For batch correction, as in previous studies Tran et al. [39] and Li et al. [40], we conducted K-means clustering within each cell type, using batch labels as ground truth, and evaluated batch mixing performance using 1-metric. In batch correction experiments, higher clustering metrics (ARI, AMI, and ASW) indicate that different batches are separate. Consequently, higher 1-metrics (1-ARI, 1-AMI, and 1-ASW) indicate that different batches of same cell type are merged instead of being separated. Results in Fig. 5f show that predicted proteins demonstrate superior batch correction performance, as evidenced by higher 1-metrics on batch level. Overall, the results in Fig. 5e and Fig. 5f indicate that the predicted proteins achieve a remarkable balance between correcting batch effects and preserving cellular heterogeneity compared to the ground truth.

All analyses above were conducted with access to the true protein value. However, scTranslator also has valuable application scenarios in the absence of fine-tuning, where only transcriptomics data is available. In this zero-shot scenario, pre-trained models can be used to directly predict proteins and potentially support downstream tasks. We utilized pre-trained scTranslator to predict 14,000 proteins, including both membrane surface and interior proteins, for each cell based on a pan-cancer scRNA-seq dataset. To demonstrate the effectiveness of these predicted protein levels, we implemented a comparison experiment between single-omic RNA and multi-omics, composed of predicted proteins and RNA. The comparison was conducted on the task of identifying the origin of the cell as a tumor or normal site. Five groups of high-variable RNA and protein were selected, ranging from 1000 to 5000 (See method section for detail). The results of these experiments are presented in Fig. 5f, which shows that the performance of the multi-omics data was superior to that of single-omic RNA in terms of accuracy, f1 score, and recall. This further demonstrates that scTranslator-generated multi-omics data can furnish more extensive insights than single-omic data. This capability can expedite a more profound comprehension of the interplay among different omics data in cellular processes and disease mechanisms.

## 4 Discussion

scTranslator is analogous to the translation process of the central dogma, which describes the flow of genetic information from RNA to protein. In a similar vein, our model bridges the gap between gene expression and protein abundance using computational methods. Therefore, it has immense potential to enhance our comprehension of cellular processes at the molecular level. By adopting customized Transformer encoder-decoder architecture, scTranslator becomes a promising approach to predict proteome abundance from single-cell transcriptome data. The model was extensively trained on large-scale datasets and achieved high accuracy in predicting proteome abundance levels on both bulk and single-cell data. Benefiting from the attention mechanisms of scTranslator, we are able to capture complex relationships between transcriptome and proteome from the model interpretability perspective. Furthermore, the flexibility of scTranslator facilitates further investigation into the impact of a specific gene on a cellular level by pseudo-knockout. Additionally, paired multiomics data were generated by pre-trained scTranslator based on the pan-cancer scRNA-seq dataset, where the proteome data was experimentally unavailable, and the transcriptome data was out-of-sample. The results suggest that the generated protein abundance, in combination with measured RNA expression, outperforms single-omic RNA in identifying tumor or normal origin cells.

scTranslator has been systematically benchmarked and demonstrated to be accurate, stable, and flexible, displaying superior performance over state-of-the-art methods across various cell types and conditions. However, there are still limitations to scTranslator. Although we have collected large-scale datasets, it is noteworthy that the scale of samples and features available for scTranslator remains limited at the single-cell level. It is anticipated that the application of multi-omics profiling will facilitate the generation of a substantial number of paired datasets. Additionally, during the preparation of this manuscript, spatial CITE-seq data is available. The current version of scTranslator does not incorporate this dataset, which is considered in the next iteration of the tool’s release. Although single-cell sequencing has the potential to enhance our understanding of cancer, which is caused by the abnormal proliferation and transformation of individual cells, the current version of scTranslator application only focuses on the cell level. Future work should aim to investigate the mechanisms of individual cells at the patient level.

Next, we will discuss the potential applications of scTranslator. Firstly, scTranslator can generate paired multi-omics data at the single-cell level. Given the abundance of single-cell transcriptomics data, scTranslator can be used to complete missing proteomics data, thereby creating multi-omics datasets. Moreover, we have released the first stage pre-trained scTranslator, which was trained on bulk datasets and can produce bulk-level multi-omics data. Secondly, the encoder of scTranslator has been pre-trained on a substantial amount of transcriptomic data and can serve as a transcriptomic embedding tool for downstream single-cell analyses, including cell cycle, cell type, and cell origin. Thirdly, we provide a demo in this manuscript that showcases the attention matrix generation process. It allows users to explore the relationship between transcriptomics and proteomics, gene regulation, and protein interactions as desired. Finally, the proposed pseudo-knockout method allows researchers to investigate the impact of genes of interest on the proteome, minimizing experimental costs before proceeding to formal wet experiments.

Overall, scTranslator represents a significant advancement in the field of single-cell multi-omics. We believe that scTranslator will be a valuable and potent tool for researchers seeking to elucidate the cellular relationship of complex biological processes. We also hope that scTranslator can open up new opportunities for comprehending a wide range of diseases, particularly those related to cancer.

## 5 Methods

### 5.1 scTranslator framework

scTranslator was developed utilizing the Transformer architecture incorporating customized design to effectively utilize all information provided by RNA and protein. Although scTranslator follows an encoder-to-decoder sequence transduction framework similar to Transformer, it does not employ masked multi-head attention in the decoder, which is used to focus information from the output embedding in an auto-regressive manner to prevent information disclosure of future positions [11]. This exclusion is based on the biological rationale that the absolute position relationship between tokens in NLP is meaningless for protein abundance data. To enable the model to differentiate between expression values from different genes, we proposed a re-indexed GPE inspired by the positional encoding from Transformer via injecting gene information into the model. To handle the challenge of extremely long sequences of RNA and protein, Performer, a linear complexity version of Transformer, was utilized instead of regular full-rank attention with quadratic space and time complexity. scTranslator was constructed using an end-to-end training strategy, whereby the model is supervised by MSE loss between predicted proteins and ground truth.

#### 5.1.1 Re-indexed gene positional encoding (GPE) and expression value embedding

Previous studies in this field have limited their scope by fixing the positions of RNA and protein and considering only the expression value of the corresponding gene. However, this approach has overlooked the valuable information available from gene names and their mutual effect. To address this deficiency, we proposed a re-indexed GPE module to utilize the previously ignored information. Specifically, we re-indexed all officially certified genes (totaling 75,500, obtained from the latest NCBI database) to discrete ID values as provided in Supplementary Table 4. We refrained from using the original gene ID number provided by NCBI due to the large maximum ID (1,255,000), which can lead to high memory usage and slow training times. Our GPE module employs an embedding layer, implemented through a trainable lookup table, to transform re-indexed gene IDs into a continuous vector space. The module supports a maximum vocabulary size of 85,500, with 10,000 reserved positions for new genes or findings not covered in the current study. Together with the rest of scTranslator, GPE module is trained concurrently to acquire informative representations of all genes.

In addition to the Gene ID, there is also a challenge of how to utilize the expression value of RNA. The standard Transformer model in NLP accepts 1D sequences of tokens and maps tokens to embedding vectors through the embedding layer. In scTranslator encoder, the expression of each gene is treated as a “token” and all expression values from one cell or patient form a “sentence”. The raw value is projected to a pre-set dimension vector by a linear embedding layer.

The incorporation of gene position embedding and expression value embedding, as illustrated in Extended Data Fig. A7, provides the potential for a comprehensive understanding of gene expression and mutual effects. Moreover, the GPE module enables scTranslator to be an align-free model, enhancing its capabilities.

#### 5.1.2 Model backbone

We adopted the Transformer encoder-decoder architecture with a multi-head FAVOR+ mechanism as the model backbone. The encoder comprises *N* = 2 identical layers, each including a sub-layer of multi-head FAVOR+ mechanism and a sub-layer of position-wise fully connected feed-forward network. Residual connections and layer normalization are employed around each of the sub-layers. The decoder is also composed of *N* = 2 identical layers, with the same sub-layers design as the encoder and an additional GPE module for denoting protein ID instead of RNA ID. The sub-layers, embedding layers, and GPE module produce outputs of dimension *d_model_* = 128. Given the current single-cell quantification coverage range of RNA and protein, the maximum sequence length of the encoder and decoder were set to 20,000 and 1,000, respectively. A multi-layer perceptron (MLP) was employed for sentence length conversion between the encoder and decoder. Additionally, an 8-headed attention mechanism was incorporated in each sub-layer. The entire model consisted of 117 million parameters.

The FAVOR+ mechanism, proposed in Performer [25], has been demonstrated to reduce the quadratic computational complexity required by regular attention mechanisms to linear complexity, both theoretically and experimentally. We assume that *X ∈ ℝ^L×dmodel^* is the input embedding including position embedding and expression value embedding with the sequence length *L*. The queries, keys, and values are intermediate representations of the input, obtained by *Q* = *W_Q_X*, *K* = *W_K_X*, *V* = *W_V_ X*. The regular bidirectional dot-product attention has the following form, where *A ∈ ℝ^L×L^* is attention matrix:

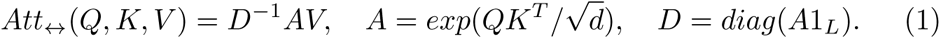

The FAVOR+ mechanism tries to find a generalized kernel to map *Q*/*K* to *Q^′^*/*K^′^* so that attention matrix *A* approximates the product of *Q^′^* and *K^′^*:

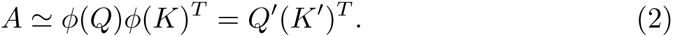

The generalized form of the mapping function is:

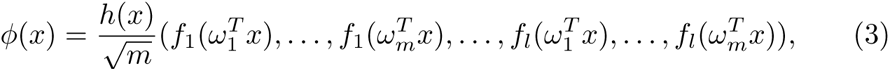

where, function *f_j_*: ℝ *→* ℝ, *j* = 1, 2*,…, l*, function *h*: ℝ*^d^ →* ℝ, and random samples: 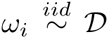 for some distribution 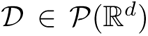. Therefore, the efficient approximate attention for the Performer is:

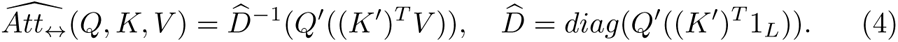

### 5.2 Datasets

As there are three stages in our framework, supervised learning on paired bulk data, supervised learning on paired single-cell data, and inference on single-cell RNA dataset with or without fine-tuning utilizing pre-trained scTranslator. In the first stage, large amounts of paired RNA and protein data were used for general pattern learning at the bulk level. In the second stage, scTranslator continues end-to-end learning on paired single-cell data. In the third stage, pre-trained scTranslator can be used to predict protein abundance in a new single-cell dataset. If there are paired RNA and protein measurements, the model can be fine-tuned using a subset of the data, and the remaining data can be used to predict the proteome based on the transcriptome. If protein data is unavailable, the model can predict protein abundance directly from the transcriptome without fine-tuning.

#### 5.2.1 Bulk datasets

The Cancer Genome Atlas (TCGA) data [41–44] in this study includes 63 datasets, consisting of all available samples with both RNA and protein simultaneously across 30 cancer types. The data from Clinical Proteomic Tumor Analysis Consortium (CPTAC) includes 7 datasets [45–51] involving endometrial carcinoma, pancreatic ductal adenocarcinoma, lung squamous cell carcinoma, lung adenocarcinoma, glioblastoma, pediatric brain cancer, and breast cancer. The dataset from Broad Institute [52, 53] is about over 1,000 cancer cell lines from individuals of various lineages and ethnicities. The dataset from Memorial Sloan Kettering Cancer Center (MSKCC) [54] is about bladder cancer. The union of these datasets involves 31 cancer types and 18,227 samples in total. In this study, we used 90% bulk samples for first-stage pre-training and the rest 10% data for evaluation.

#### 5.2.2 Single-cell datasets

##### The Seurat v4 PBMCs dataset

The Seurat v4 PBMCs dataset, provided by reference [24], integrated 161,764 human peripheral blood mononuclear cells (PBMCs) with panels extending to 224 antibodies measured by CITE-seq. The number of transcriptome genes is 23,385. In this study, we used 90% of the Seurat v4 PBMCs dataset for the second-stage pre-training and the rest for performance testing as Dataset 1.

##### The REAP-seq PBMCs dataset

The REAP-seq PBMCs dataset, published in reference [55], contains 4,330 PBMCs with simultaneous measurements of 44 proteins and 21,005 transcriptome genes. In this study, we used the REAP-seq PBMCs dataset for the third-stage inference as Dataset 2.

##### The CITE-seq CBMCs dataset

The CITE-seq CBMCs dataset, published in reference [38], provided simultaneous measurements of 13 cellular proteins and 16,508 transcriptome genes. This dataset includes 8,005 cord blood mononuclear cells (CBMCs). In this study, we used the CITE-seq CBMCs dataset for the third-stage inference as Dataset 3.

##### The single-cell pan-cancer dataset

The single-cell pan-cancer dataset, provided by reference [56], integrated tumor-infiltrating myeloid cells from 43 patients with eight types of cancer. This dataset covers 65,698 myeloid cells with only single-cell transcriptome data, involving 15,844 genes. In this study, we utilized the single-cell pan-cancer dataset for the third-stage protein inference, followed by a cell origin exploration based on multi-omics analysis.

### 5.3 Data pre-processing

In light of the fact that scTranslator has a capacity of 20,000 genes as input, to reserve full gene-level information and interpretability, we applied no dimension reduction, either by feature selection or feature extraction. As for the count matrix of RNA or protein, scaled Max-Min normalization [57] was performed on data across each cell, formalized as:

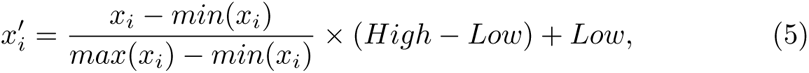

where, cell index: *i* = 1, 2*,…, num_cell_*, scale factor: *High* = 1 and *Low* = 10*^−^*^8^. The purpose of the scale factor is to distinguish between the genuine count of zero, which is scaled to 10*^−^*^8^ in equation (5), and the padding of zero used for the model input.

### 5.4 Systematic benchmark implementation

The cTP-net and sciPENN were executed using the Python script provided at https://github.com/zhouzilu/ctpnetpy/tree/master/ctpnet and https://github.com/jlakkis/sciPENN, respectively. For each method, the officially recommended hyper-parameter settings were adopted. We compared the performance of scTranslator and SOTA on Dataset 1, 2, and 3 under aligned, unaligned, and few-shot mode, where 9 groups of comparison in total. For both the aligned and unaligned experiments, we utilized 90% of the data for training and reserved the remaining 10% for testing. For the few-shot experiment, only 20 cells were used for training and the remaining cells were used for testing. To ensure the reliability and reproducibility of our results, we repeated each group of comparison experiments 10 times using diverse random seeds to partition the training and test data. By repeating the experiments with different seeds, we can get a better understanding of the range of possible outcomes and increase our confidence in the generalization of our findings. To ensure an equitable comparison, all models were constructed utilizing the identical data partitioning scheme. All experiments were conducted on the same hardware, consisting of eight V100 GPUs.

MSE and cosine similarity were used to evaluate the performance of models on protein inference. Supposing that the ground truth/prediction of protein is *Y* /*Ŷ ∈* ℝ*^n^*, where *n* is the number of proteins and *y_i_*/*ŷ_i_* represents the ground truth/prediction of *i_th_* protein, the MSE is then defined as follows:

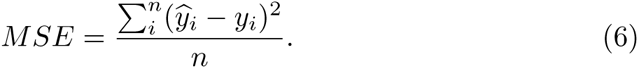

The cosine similarity can be described as:

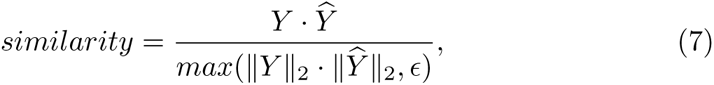

where *ɛ* = 1*e^−^*^8^ is a small value to avoid division by zero.

### 5.5 Attention score based regulatory inference

The attention mechanism can identify important features and interactions among data points, making it well-suited for regulatory inference in gene expression analysis. In this paper, we implemented a comprehensive interpretability analysis to identify important genes and interactions between RNA and proteins.

The score in the attention matrix represents the relevance or importance of a specific part from the input sequence to a given task or output. In scTranslator, the attention score in the encoder reflects the contribution of each RNA and the interaction of RNA pairs. Similarly, we can identify the key protein and protein–protein interactions by the attention matrix of decoder. What’s more, the attention matrix from encoder to decoder revealed the importance of RNA when inferring protein. The attention matrix can be obtained from equation 4, which entails taking the dot product between the queries and keys, while also replacing *V* with a diagonal matrix. We calculate the attention matrix for all the samples and take their sum as the final attention matrix. Take the encoder-to-decoder attention matrix as an example, in this attention matrix, each value *A*(*i, j*) represented how much attention from protein *i* was paid to gene *j*. The attention matrix is visualized by heatmap. Furthermore, we implemented enrichment analysis on top-ranked genes using the method on the g:Profiler [58] website https://biit.cs.ut.ee/gprofiler.

### 5.6 Pseudo-knockout

Gene knockout experiments are an essential tool for understanding gene function and its contribution to biological processes. By knock-down or knock-out specific genes, researchers can investigate the effects on cellular and organismal phenotypes, providing insights into the gene’s role in development, disease, and other biological pathways. Gene knockout experiments also enable the validation of potential therapeutic targets and the development of new treatments for genetic disorders. However, traditional wet-lab gene knockout experiments can be time-consuming and expensive, making it difficult to explore the full range of possibilities.

In this study, we employed the pre-trained scTranslator to investigate the impact of RNA gene on proteins by pseudo-knockout, encompassing both surface and intracellular proteins. This pseudo-knockout was conducted by removing a specific gene in RNA and comparing the change in protein abundance before and after the knockout of this gene. In this paper, we explored the impact of removing TRIM39, MTHFR, AKT1, ESR1, IL6, VEGFA, EGFR, TNF, and TP53. These pseudo-knockout experiments feed single-cell RNA data of Dataset 1 and query ID of 14,000 proteins that the model learned before. Finally, we reported the top influenced proteins.

### 5.7 Cell marker analysis and cell clustering

For cell marker analysis, we utilized the Python package “Scanpy” (v.1.9.1). The cell type marker is obtained through “*scanpy.tl.rank genes groups*” by differential gene expression analysis based on their statistical significance and fold change between different cell types. To visualize the marker across different cell types, UMAP is plotted through “*scanpy.pl.umap*” under 50 neighborhoods. The predicted protein, true protein, and true RNA adopted the same parameter setting in this part.

Cell clustering is conducted by “*sklearn.cluster.KMeans*”. The number of clusters to form as well as the number of centroids to generate in K-means is equal to the ground truth of level-1 cell types in Dataset 1. Cell clustering performance of each omic data was evaluated by ARI, AMI, and ASW provided by scikit-learn (v.1.1.2) at https://scikit-learn.org/stable/modules/clustering. html#clustering-performance-evaluation.

### 5.8 Cell origin analysis based on pan-cancer data

In this experiment, we utilized the pre-trained scTranslator to predict protein abundance based on RNA data from the single-cell pan-cancer dataset and query ID of 14,000 proteins. Subsequently, we developed a cell origin classifier, similar to the encoder of scTranslator with N = 1 identical layers and *d_model_* = 32, to distinguish cells derived from tumor tissue versus normal tissue. To investigate whether predicted protein abundance could aid in cell origin identification, we compared single-omic RNA data with multi-omics data consisting of RNA and pseudo-quantified proteins, using the same cell origin classifier. To minimize the effect of chance, we selected highly variable RNA and protein genes through “*scanpy.tl.rank genes groups*”. Five groups of RNA and proteins were selected, where the number of genes changes from 1,000, 2,000, 3,000, 4,000, to 5,000. The max seq length of this cell origin classifier is determined by the number of genes. We reported the mean and variance of accuracy, f1 score, and recall [57] in all 5 runs.

## 6 Data availability

All datasets used in this study were obtained from public data repositories. The TCGA data was downloaded from https://www.cancer.gov/ ccg/research/genome-sequencing/tcga. The CPTAC data was downloaded from https://www.cbioportal.org/study/summary?id=ucec cptac 2020, https://www.cbioportal.org/study/summary?id=paad cptac 2021, https://www.cbioportal.org/study/summary?id=lusc cptac 2021, https://www.cbioportal.org/study/summary?id=luad cptac 2020, https://www. cbioportal.org/study/summary?id=gbm cptac 2021, https://www.cbioportal. org/study/summary?id=brain cptac 2020, https://www.cbioportal.org/ study/summary?id=brca cptac 2020. The Broad Institute CCLE data was downloaded from https://sites.broadinstitute.org/ccle. The MSKCC data was downloaded from https://www.cbioportal.org/study/summary? id=blca msk tcga 2020. The Seurat v4 PBMCs dataset was downloaded from https://www.ncbi.nlm.nih.gov/geo/query/acc.cgi?acc=GSE164378 and https://atlas.fredhutch.org/nygc/multimodal-pbmc/. The CITE-seq CBMCs dataset was downloaded from https://www.ncbi.nlm.nih.gov/geo/query/ acc.cgi?acc=GSE100866. The REAP-seq PBMCs dataset was downloaded from https://www.ncbi.nlm.nih.gov/geo/query/acc.cgi?acc=GSE100501. The single-cell pan-cancer dataset was downloaded from https://www.ncbi.nlm.nih.gov/geo/query/acc.cgi?acc=GSE154763.

## 7 Code availability

scTranslator is available at https://github.com/TencentAILabHealthcare/ scTranslator. For reproducibility, the scripts for all experiments and results analysis were also included in the above repository.

## 8 Author contributions

L.L., F.Y., and J.Y. conceived the project. L.L. designed the algorithm framework and developed the method. L.L. performed research and conducted experiments under the supervision of K.W., F.Y., and J.Y.. L.L. and F.Y. analysed the results and wrote the manuscript. K.W. and J.Y. revised the manuscript. W.L. provided suggestions for downstream analysis, assisted with figure polishing, and contributed to manuscript improvement. All authors reviewed and approved the manuscript.

## Supporting information

Supplementary Table 1 Protein predictability and functional enrichment results.xlsx

Supplementary Table 2 Systematic benchmark results.xlsx

Supplementary Table 3 Reported and novel markers.xlsx

Supplementary Table 4 Re-indexed Gene ID.xlsx

## Acknowledgements

We thank Zetian Zheng, Duyu Tang and Fang Wang for their valuable suggestions and discussion during the preparation of this manuscript. This research was substantially sponsored by the research projects (Grant No. 32170654 and Grant No. 32000464) supported by the National Natural Science Foundation of China and was substantially supported by the Shenzhen Research Institute, City University of Hong Kong. The work described in this paper was substantially supported by the grant from the Research Grants Council of the Hong Kong Special Administrative Region [CityU 11203723]. This project was substantially funded by the Strategic Interdisciplinary Research Grant of City University of Hong Kong (Project No. 2021SIRG036). The work described in this paper was partially supported by the grant from City University of Hong Kong (CityU 9667265) and Key-Area Research and Development Program of Guangdong Province (2021B0101420005).

## Appendix A Extended Data

**Fig. A1.**
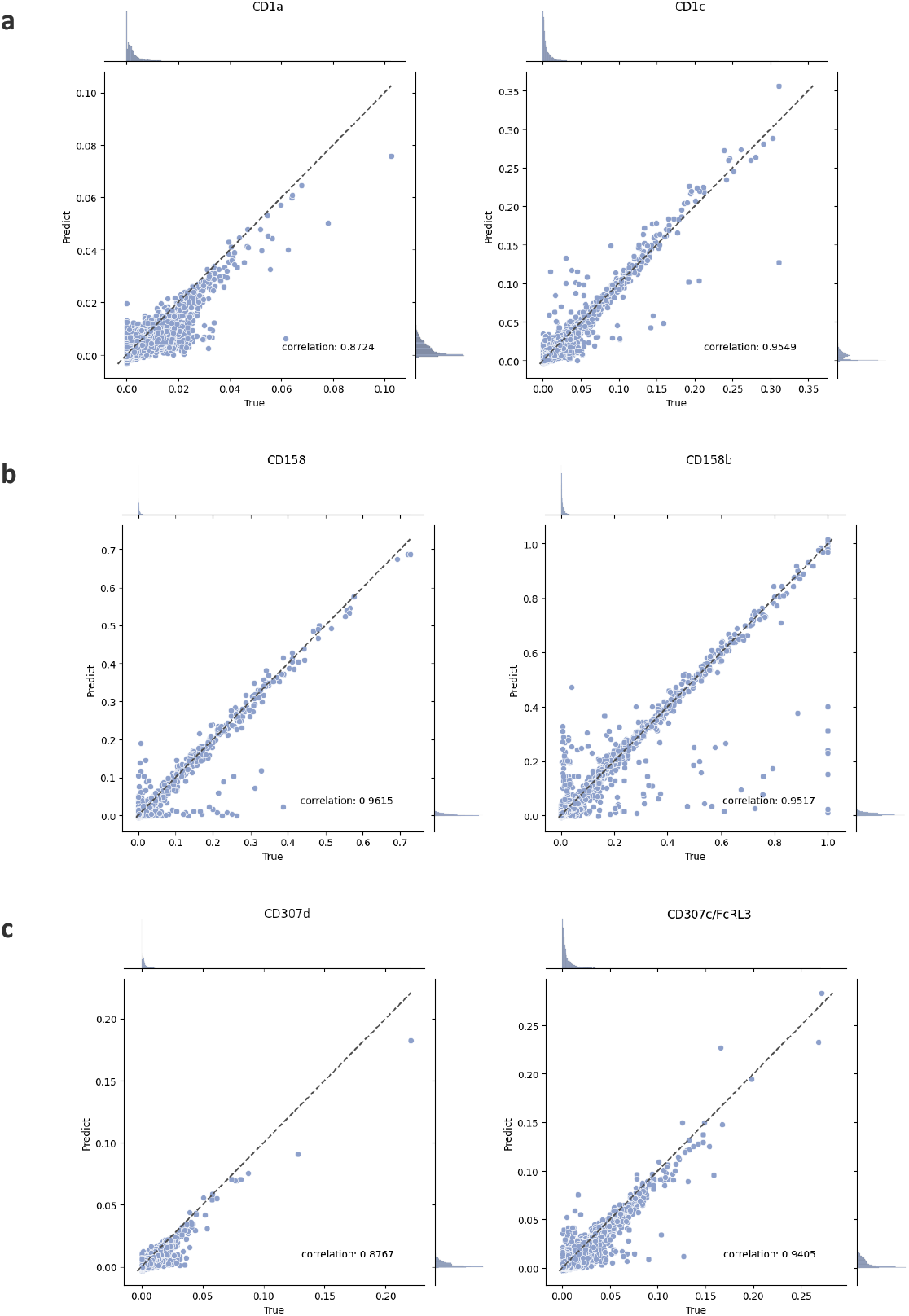
Joint plots for evaluating prediction performance at individual level. **a-c** Joint plots of distribution histogram and scatter plot for CD1, CD158 and CD307 protein family members.

**Fig. A2.**
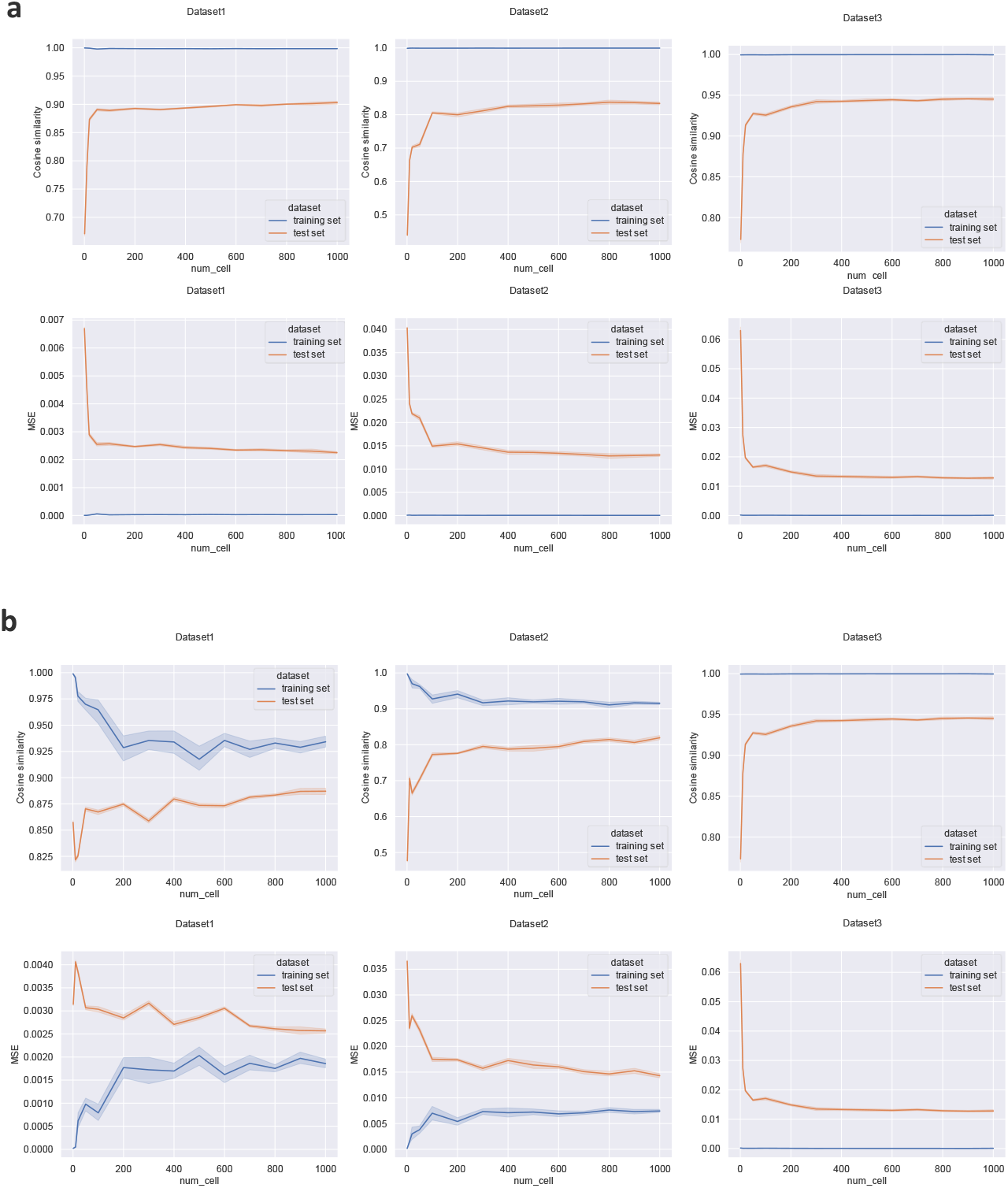
Influence of sample amount on scTranslator in fine-tuning stage. **a,** Experiment results of sample amount effect for aligned mode. **b,** Experiment results of sample amount effect for un-aligned mode.

**Fig. A3.**
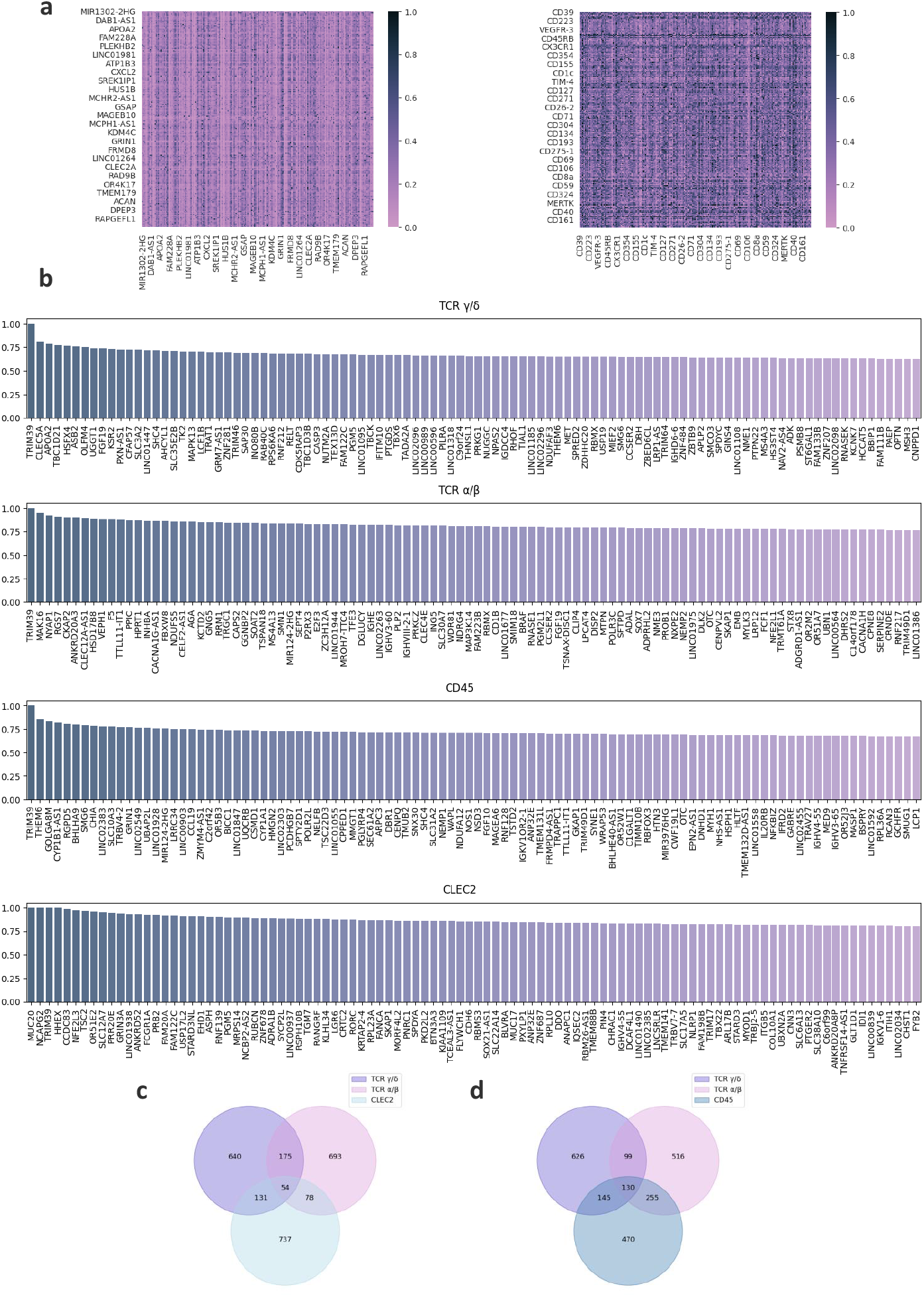
Gene regulatory and protein interaction investigate. **a,** Heatmap of attention score matrices, where the left and right panels correspond to the encoder’s and decoder’s attention scores, respectively. **b,** Bar plot showing the most highly focused gene for proteins of TCR *α*/*β*, TCR *γ*/*δ*, CD45, and CLEC2. **c, d** The Venn diagram of the most focused top 5% gene intersections. **c** displays TCR *α*/*β*, TCR *γ*/*δ*, and CLEC2, whereas the **d** exhibits TCR *α*/*β*, TCR *γ*/*δ*, and CD45.

**Fig. A4.**
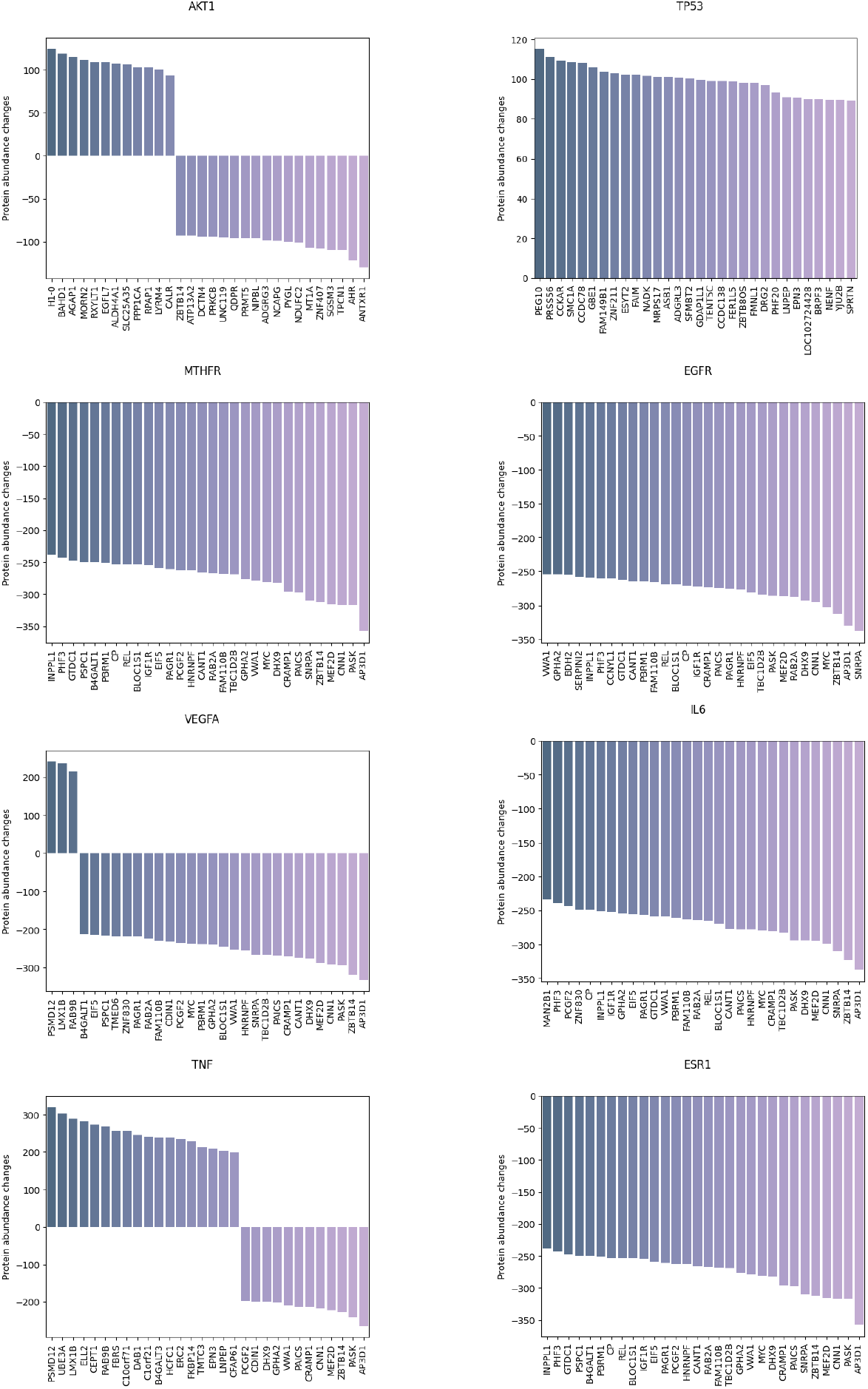
Pseudo-knockout results of well-studied genes. The bar chart shows the most variable proteins affected by knocked gene perturbation. The title of each panel denotes the knocked-out gene.

**Fig. A5.**
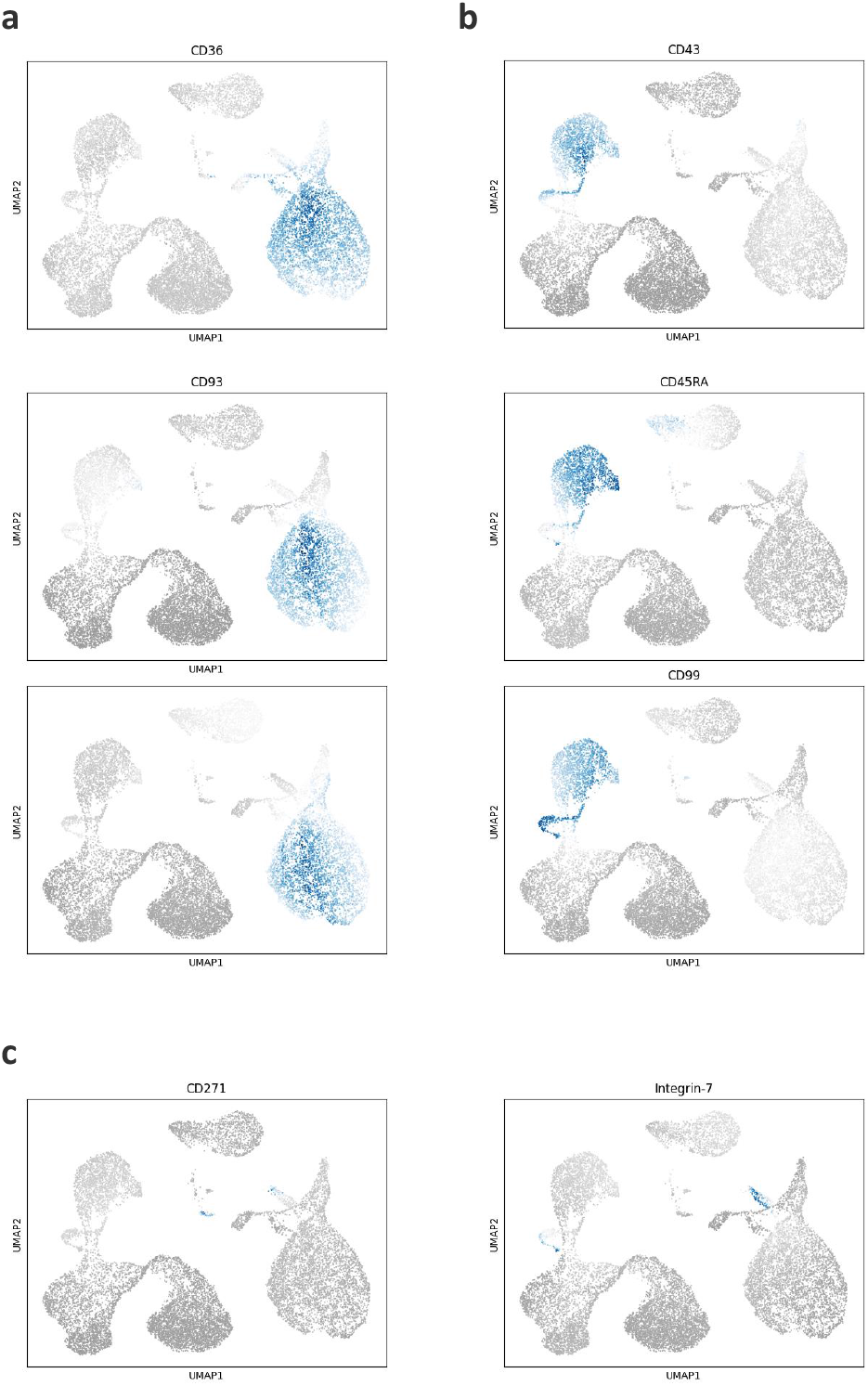
UMAP visualization of novel markers. **a,** UMAP plots of predicted protein colored by expression level of three new markers for monocyte. **b,** UMAP plots of predicted protein colored by expression level of three new markers for NK cells. **c,** UMAP plots of predicted protein colored by expression level of two new markers for DC and its subtype.

**Fig. A6.**
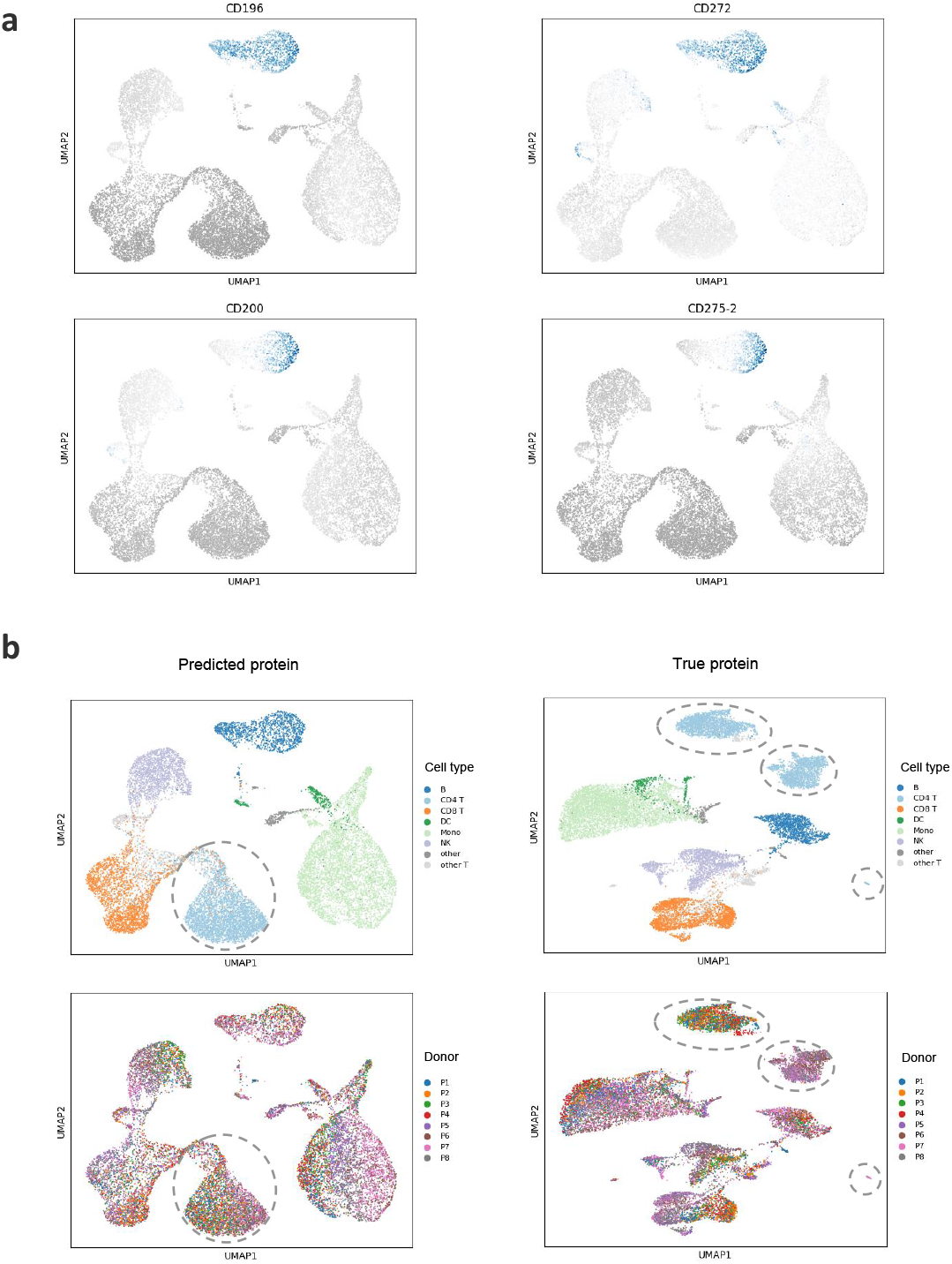
UMAP visualization to observe batch effect and novel marker. **a,** UMAP plots of predicted protein colored by expression level of four new markers for B cell and its subtypes. **b,** UMAP plots of predicted protein on Dataset 1, colored by cell type in top panel and by batch information in bottom panel. The left plots are visualization for inferred protein by scTranslator and the right plots are visualization for true protein.

**Fig. A7.**
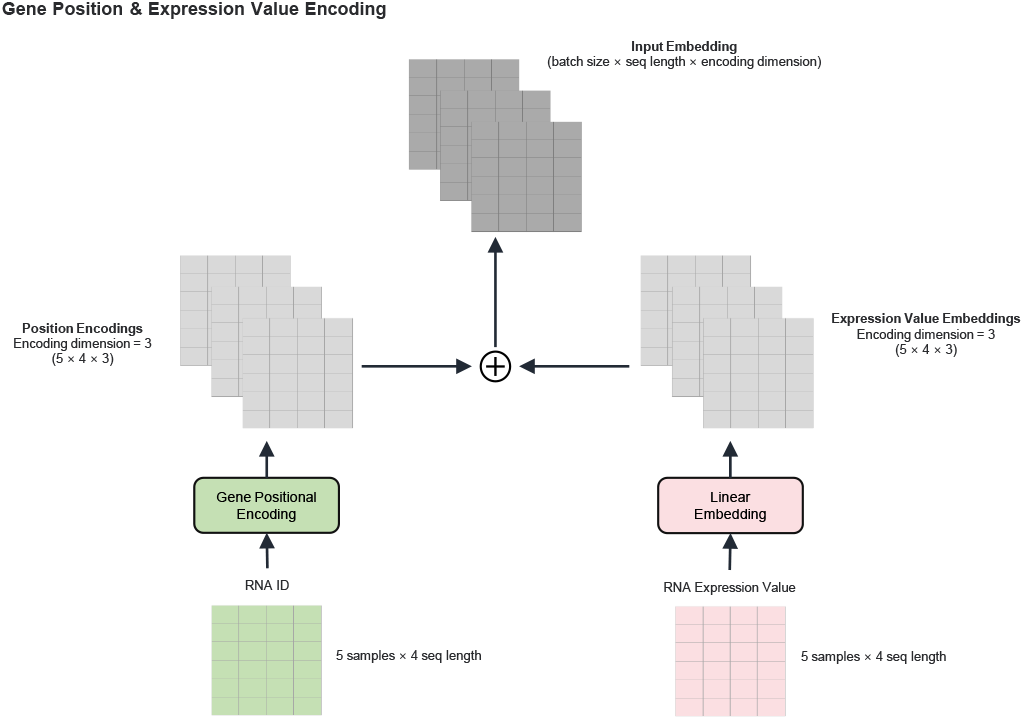
Illustration of gene position embedding and expression value embedding.

## Supplementary information

Supplementary Table 1: Predictability and functional enrichment results of proteins.

Supplementary Table 2: Systematic benchmark results.

Supplementary Table 3: Reported and novel markers. Supplementary Table 4: Re-index Gene ID

